# Neural gain control measured through cortical gamma oscillations is associated with individual variations in sensory sensitivity

**DOI:** 10.1101/348656

**Authors:** EV Orekhova, TA Stroganova, JF Schneiderman, S Lundström, B Riaz, D Sarovic, OV Sysoeva, C Gillberg, N Hadjikhani

**Affiliations:** University of Gothenburg, Gillberg Neuropsychiatry Centre (GNC), Gothenburg, Sweden; Moscow State University of Psychology and Education, Center for Neurocognitive Research (MEG Center), Moscow, Russia; Moscow State University of Psychology and Education, Autism Research Laboratory, Moscow, Russia; University of Gothenburg, Institute of Neuroscience &Physiology, Department of Clinical Neurophysiology and MedTech West, Gothenburg, Sweden; Harvard Medical School, MGH/MIT/HST Martinos Center for Biomedical Imaging, Charlestown, MA USA

**Keywords:** Gamma oscillations, response gain control, sensory sensitivity, visual motion, magneto-encephalography (MEG), autism spectrum disorders (ASD)

## Abstract

Gamma oscillations facilitate information processing by shaping the excitatory input/output of neuronal populations, and their suppression by strong excitatory drive may stem from inhibitory-based gain control of network excitation. Individual variations in the gamma suppression may therefore reflect efficiency of gain control and subjective sensitivity to everyday sensory events. To test this prediction, we assessed the link between self-reported sensory sensitivity and changes in magneto-encephalographic gamma oscillations as a function of motion velocity of high-contrast visual gratings. The induced gamma oscillations increased in frequency and decreased in power with increasing stimulation intensity. As expected, weaker suppression of the gamma response correlated with sensory hypersensitivity. Robustness of this result was confirmed by its replication in the two samples: neurotypical subjects and people with autism, who had generally higher sensory sensitivity. We conclude that intensity-related suppression of gamma response is a promising biomarker of homeostatic control of the excitation-inhibition balance in the visual cortex.

## Introduction

The balance between excitation and inhibition (E/I balance) in neural networks orchestrates neural activity in space and time, and is important for cortical functioning (Dorrn et al., 2010; Isaacson and Scanziani, 2011; Xue et al., 2014). Activity of the excitatory (E) and inhibitory (I) neurons is fine-balanced in the normal brain and this balance is disrupted in epilepsy (Dehghani et al., 2016) and neurodevelopmental disorders, such as e.g. autism and schizophrenia (LeBlanc and Fagiolini, 2011; Lee et al., 2017; Nelson and Valakh, 2015; Rubenstein and Merzenich, 2003).

Animal findings promote the discoveries of new drugs aimed to restore the neural E/I balance in the patients with brain and mental health disorders (Lee et al., 2017; Tu et al., 2017), however there are still considerable challenges in testing them in clinical trials. A lack of quantifiable non-invasive measures of the E/I balance in the human brain precludes stratification of heterogeneous patient populations according to the distinct E/I balance subtypes, and hinders assessment of the treatment efficacy (Ecker et al., 2013; Levin and Nelson, 2015).

The stimulus-induced high-frequency MEG/EEG gamma oscillations (30-100 Hz) have attracted considerable attention as a putative noninvasive indicator of an altered E/I balance in human cortex (Levin and Nelson, 2015; Nelson and Valakh, 2015). Gamma oscillations are generated by populations of interconnected excitatory and inhibitory neurons and are intimately related to the balance between inhibitory and excitatory neurotransmission (Buzsaki and Wang, 2012; Vinck et al., 2013). However, the current attempts to define a single parameter of human gamma response that would accurately capture the E/I balance have led to ambiguous results (Cousijn et al., 2014; Edden et al., 2009; Perry et al., 2014).

In a previous study we put forward the idea that it is possible to estimate efficiency of the E/I balance regulation in the visual cortex through probing input-output gain in the strength of the visual gamma oscillations recorded by MEG (Orekhova et al., 2018). Response gain control is a basic property of neural networks that works to both amplify the neuronal responses to weak sensory signals and to saturate/suppress these responses under conditions of excessive input (e.g. (Peirce, 2007)). The strength of visually induced gamma oscillations is controlled in accord with this mechanism, in both animals (Jia et al., 2011; Jia et al., 2013; Roberts et al., 2013; Salelkar et al., 2018) and humans (Orekhova et al., 2018), wherein a gradual increase in excitatory drive elicits an increase in gamma power up to a certain ′transition′ point, and a stimulation intensity past that point leads to suppression of the gamma response.

According to modeling studies, suppression of the oscillatory gamma response at high intensities of excitatory drive is caused by over-excitation of the I-neurons resulting in the loss of neural synchrony (Borgers and Kopell, 2005; Borgers and Walker, 2013; Cannon et al., 2014). Computational models further suggest that the gamma suppression is substantially reduced when the excitation of the E-neurons is disproportionally higher than that of the I-neurons (i.e. in the case of a high E/I ratio) (Borgers and Kopell, 2005). Indeed, opto-genetic research in animals demonstrated that gamma oscillations are particularly powerful when the high excitation of excitatory neurons is not properly balanced by inhibition (Yizhar et al., 2011). In regard to humans, these considerations imply that the brain of individuals exhibiting weaker suppression of the visual MEG gamma response is characterized by less efficient inhibitory-based capacity to down-regulate the rising excitation, i.e. by an E/I ratio shifted towards excitation (see (Orekhova et al., 2018) for discussion).

If suppression of the visual gamma response does reflect the capacity to regulate the E/I balance in the visual cortex, this phenomenon should have behavioral manifestations. On the behavioral level, the enhanced neural excitability of sensory cortices is associated with heightened or aversive reactions to intensive sensory input. Indeed, subjective discomfort associated with intensive visual stimulation correlates positively with hemodynamic responses in the visual cortex (Bargary et al., 2015; Haigh et al., 2013). Moreover, people with neurological or neuropsychiatric disorders characterized by overt clinical symptoms of elevated neuronal excitability, such as migraine with visual aura (Boulloche et al., 2010; Maniyar et al., 2014; O’Hare and Hibbard, 2016) or epilepsy (van Campen et al., 2015) often suffer from sensory hypersensitivity. It seems plausible that the atypically strong cortical responses observed in people reporting sensory hypersensitivity are caused by deficiency of the gain control mechanisms that balance excitation and inhibition in the sensory cortices. Therefore, we predicted that reduced gamma suppression at high intensities of visual input would be associated with enhanced sensory sensitivity in everyday life.

Here, we sought to test this prediction in two independent samples of subjects: neurotypical individuals (NT) and high-functioning individuals with autism spectrum disorder (ASD). A large proportion of people with ASD are hypersensitive to environmental stimuli of different modalities. Considerable variations in sensitivity to sensory events are also present in the general population (Horder et al., 2014; Little et al., 2017), correlate with autistic features (Horder et al., 2014; Robertson and Simmons, 2013), and share with them a common genetic basis (Taylor et al., 2018). Therefore, we expected that the similar neuro-behavioral association should characterize both groups. To test this prediction, we assessed behavioral sensory sensitivity using the Adolescent/Adult Sensory Profile (A/ASP) questionnaire (Brown and Dunn, 2002) and measured velocity-related suppression the visual MEG gamma response in adults with and without ASD. Since we used moving visual stimuli, we expected to find the most prominent neurobehavioral correlations for the visual modality, and particularly for sensitivity to visual motion.

## Results

### Adolescent/Adult Sensory Profile (A/ASP)

To measure subjective sensory sensitivity we used the ‘Sensory Sensitivity’ scale of the A/ASP questionnaire that characterizes noticing behaviors, distractibility, and discomfort with sensory stimuli of different modalities. Since we measured the gamma response to moving visual stimuli, we also assessed discomfort associated specifically with intensive visual motion by combining two items of the A/ASP into the ‘Visual Motion Sensitivity’ scale (see Materials and methods for details). Additionally, we calculated the ‘Visual Low Thresholds’ – the A/ASP scale that measures a person’s notice of, or annoyance with different types of visual stimuli.

The general Sensory Sensitivity was marginally higher in the ASD than in NT participants (T_(37)_=2.25, p=0.03), while no group differences were found for the visual scales (Table 1). In *Supplementary Table 1*, we present the results of group comparisons for all standard A/ASP scales.

**Table 1.** Group differences in the A/ASP measures of sensory sensitivity.

| A/ASP item | NT (19) | ASD (20) | T-statistic |
| --- | --- | --- | --- |
| Sensory Sensitivity | 31.2 | 37.7 | -2.3* |
| *A/ASP-derived measures of visual sensitivity* |  |  |  |
| Visual Low Threshold | 12.9 | 14.9 | -1.5 |
| Visual Motion Sensitivity | 4 | 4.7 | -1.0 |

\* $p<0.05$

### MEG gamma responses in NT and ASD

The experiment was designed to modulate the intensity of visual input by varying motion velocity of high-contrast annular gratings (1.2, 3.6 or 6.0%) (Fig. 1). We localized the MEG signal in the brain and found the ‘maximally induced voxel’ (Tan et al., 2016) – the source in the visual cortex that displayed the greatest gamma response to the stimulation (see Materials and methods for details).

In both groups, the average location of the maximally induced voxel corresponded to the left calcarine sulcus and did not significantly differ between groups for either x, y or z coordinates (NT: x=−0.18, y=−9.34, z=−0.15; ASD: x=0.23, y=−9.49, z=0.13 cm). Figure 2 shows the average source localization of the motion-related gamma response measured as the weighted peak power (see Materials and methods for details) in the ASD and NT participants. These results indicate that in both groups, and in all experimental conditions, the gamma response predominantly reflects activity in the primary visual cortex.

For further analysis we averaged the response spectra across the maximally induced voxel and the 25 closest voxels.

**Figure 1.**
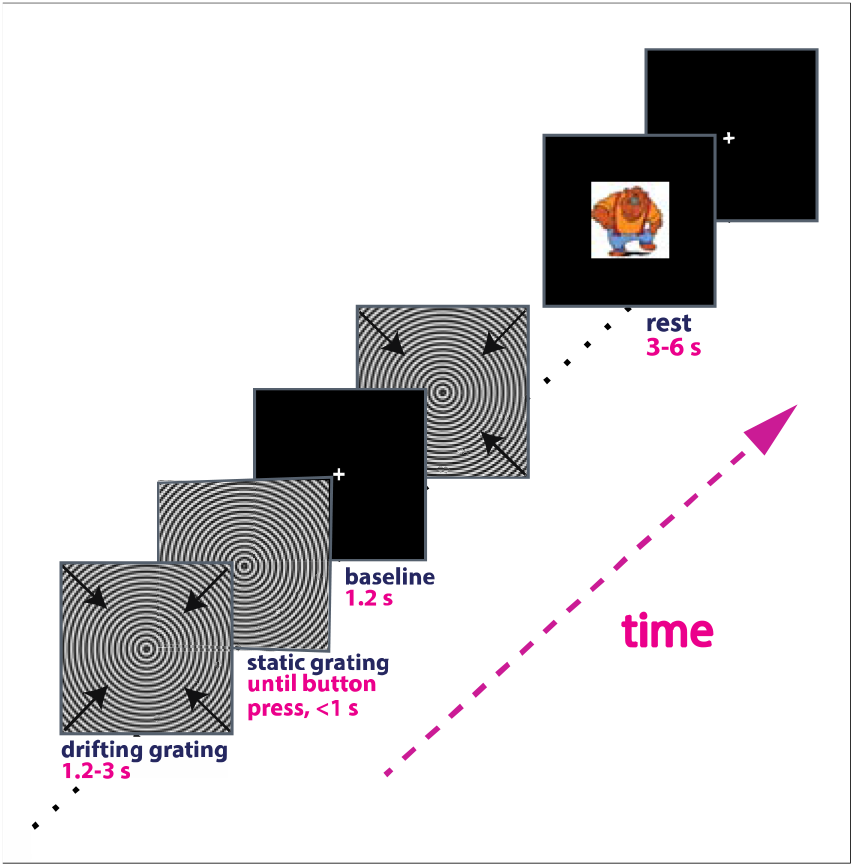
Experimental design. Each trial began with presentation of a fixation cross that was followed by an annular grating drifting inward for 1.2-3 seconds at one of the three velocities: 1.2, 3.6, 6.0°/s. Hereafter, we referred to these velocities as ‘slow’, ‘medium’, and ‘fast’. Arrows indicate direction of the motion. Participants responded to the termination of motion with a button press. Short (3-6 s) animated cartoon characters were presented randomly between every 2-5 stimuli to sustain vigilance and reduce visual fatigue. (See Materials and methods for details).

**Figure 2.**
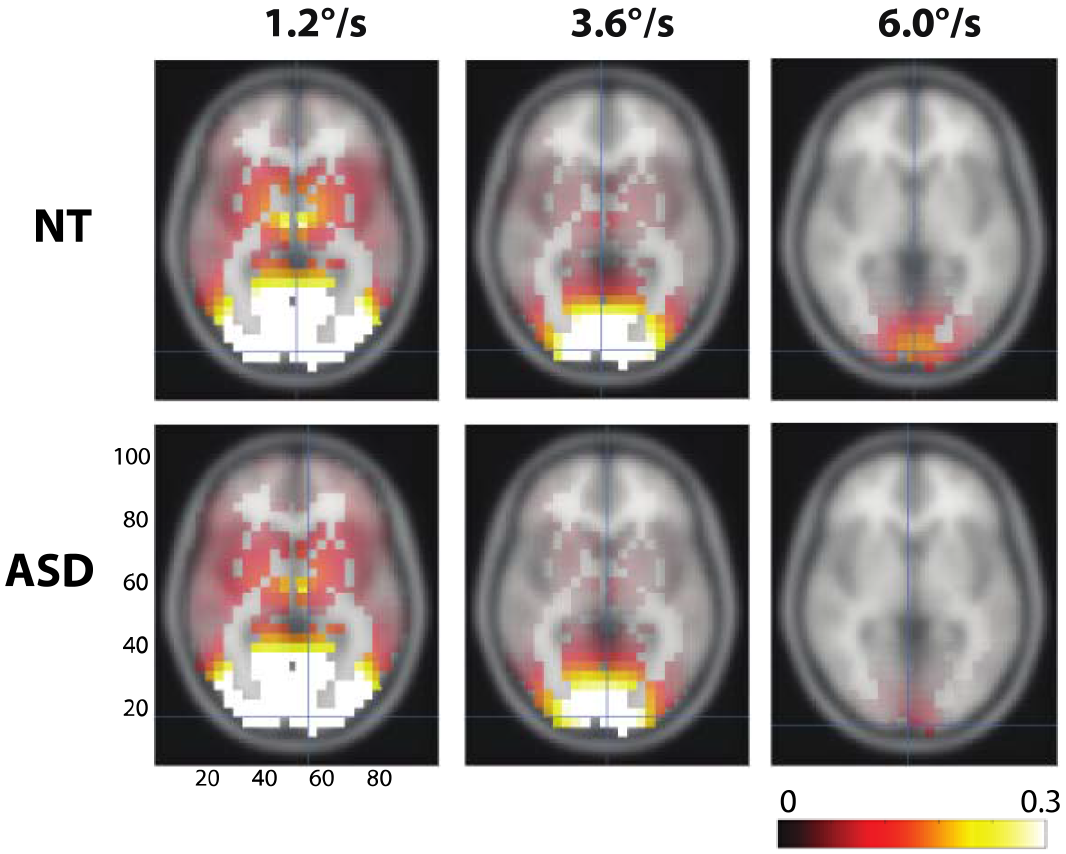
Grand average source localization of the visual gamma responses in ASD and NT individuals. Here and hereafter the magnitude of the gamma response was calculated as the ratio: (stimulus-baseline)/baseline. In both groups and under all three velocity conditions the gamma power ratio was maximal in the calcarine sulcus (marked with a blue crosshair).

Previous studies suggest that visually induced gamma oscillations might be altered in people with ASD (Dickinson et al., 2015; Stroganova et al., 2012). To test for the group differences in gamma parameters, as well as for effect of condition and its interaction with the experimental group, we used repeated measures ANOVA (rmANOVA) with the factors GROUP and VELOCITY. The power of the gamma response strongly decreased with increasing velocity (F_(2,74)_=74.2, p<0.00001), but neither effect of GROUP nor GROUP × VELOCITY interaction were significant for the gamma power (p’s>0.2) (Fig. 3 A). To analyze the frequency of the gamma response, we measured weighted gamma frequency in those subjects and conditions where the stimulus-related increases in gamma power were reliable at p<0.0001 level (see Materials and Methods for details). The rmANOVA was performed in 15 NT and 11 ASD subjects in whom the frequency was possible to assess in each of the three velocity conditions. For the gamma frequency the rmANOVA revealed highly reliable increase in frequency with increasing motion velocity (F_(2,48)_=152.6, p<0.00001), but no effect of GROUP or GROUP × VELOCITY interaction (p’s>0.5) (Fig. 3 B). To sum up, suppression of gamma response power and increase of gamma response frequency with increasing motion velocity were observed in both NT and ASD individuals and did not differ between the groups.

**Figure 3.**
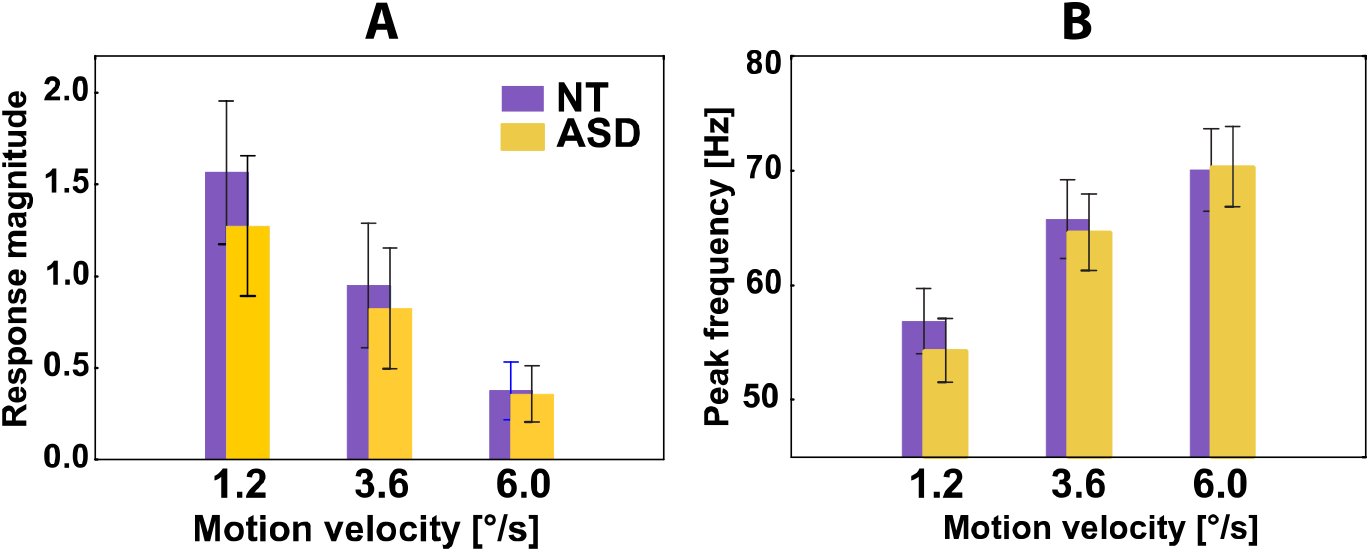
Magnitude (A) and peak frequency (B) of gamma responses to moving gratings in NT and ASD individuals. Parameters of the gamma response were measured in its focus in the visual cortex (see Materials and methods for details). Vertical bars denote 0.95 confidence intervals.

### Gamma suppression and sensory sensitivity

To quantify the velocity-related suppression of gamma response power, we introduced the ‘gamma suppression slope’ (GSS) parameter (Figure 4, see Materials and methods for details), where a more negative GSS value corresponds to stronger suppression.

**Fig. 4.**
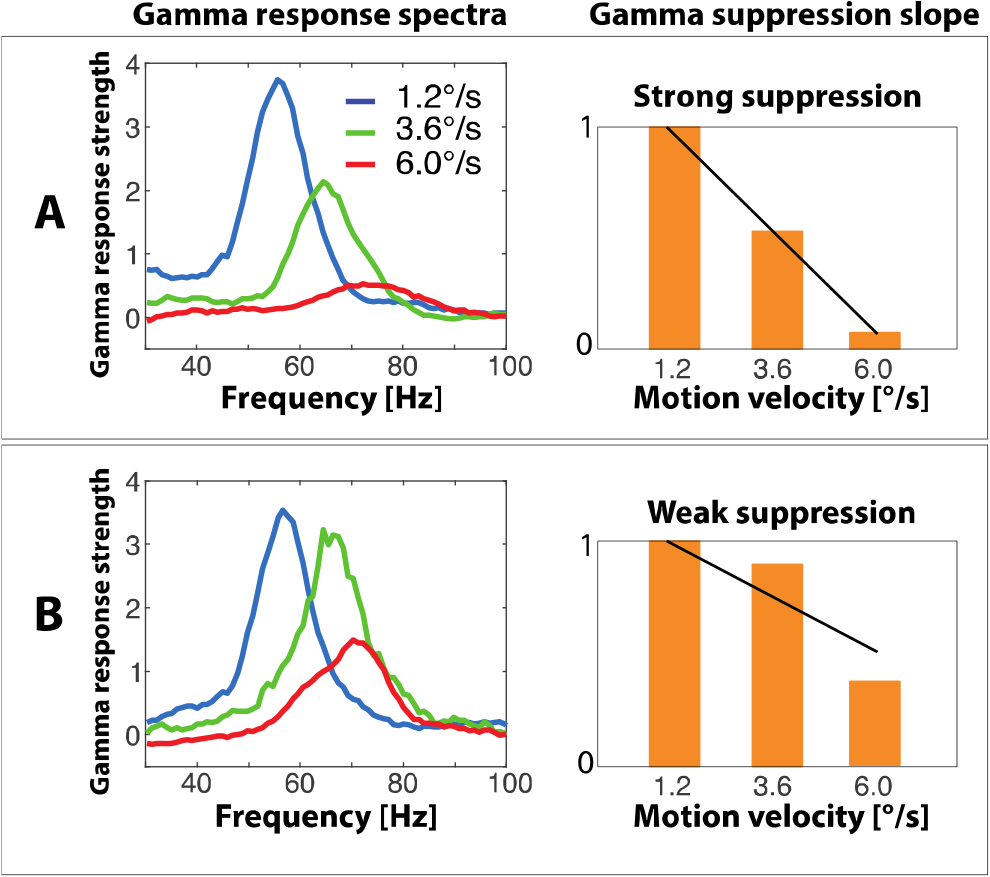
Gamma suppression slope (GSS) reflects strength of the gamma response suppression caused by increasing velocity of visual motion. Subject **A** shows a strong suppression of gamma response power with increasing motion velocity reflected in strongly negative slope of the regression line. Subject **B** has a less prominent gamma suppression corresponding to less negative GSS value.

Because the ASD and NT groups differed in neither visual sensitivity nor gamma parameters, we combined them for the correlation analysis. Table 2 and figure 5 show Spearman correlations between the sensitivity measures and the gamma suppression slope. As expected, less negative GSS (i.e. less prominent gamma suppression) correlated with higher scores on Sensory Sensitivity, visual Low Threshold, and Visual Motion Sensitivity scales. The correlation between Sensory Sensitivity and GSS remained significant when tested separately in the NT and the ASD groups. For the visual sensitivity measures the correlations with the GSS were in the same direction in the NT and the ASD groups, but did not reach significance level in the NT subjects. Correlations between the GSS and other A/ASP scales are presented in the *Supplementary Table 2* for comparison.

**Table 2.** Spearman correlations between gamma suppression slope and A/ASP sensitivity measures.

| A/ASP item | NT+ASD (38) | ASD (19) | NT (19) |
| --- | --- | --- | --- |
| *A/ASP quadrants* |  |  |  |
| Sensory Sensitivity | 0.5** | 0.57* | 0.5* |
| *A/ASP-derived measures of visual sensitivity* |  |  |  |
| Visual Low Threshold | 0.47** | 0.62** | 0.32 |
| Visual Motion Sensitivity | 0.51*** | 0.63** | 0.41<sup>#</sup> |

Significance values \* $p < 0.05$ , \*\* $p < 0.01$ , \*\*\* $p < 0.001$ , <sup>#</sup> $p < 0.1$

**Figure 5.**
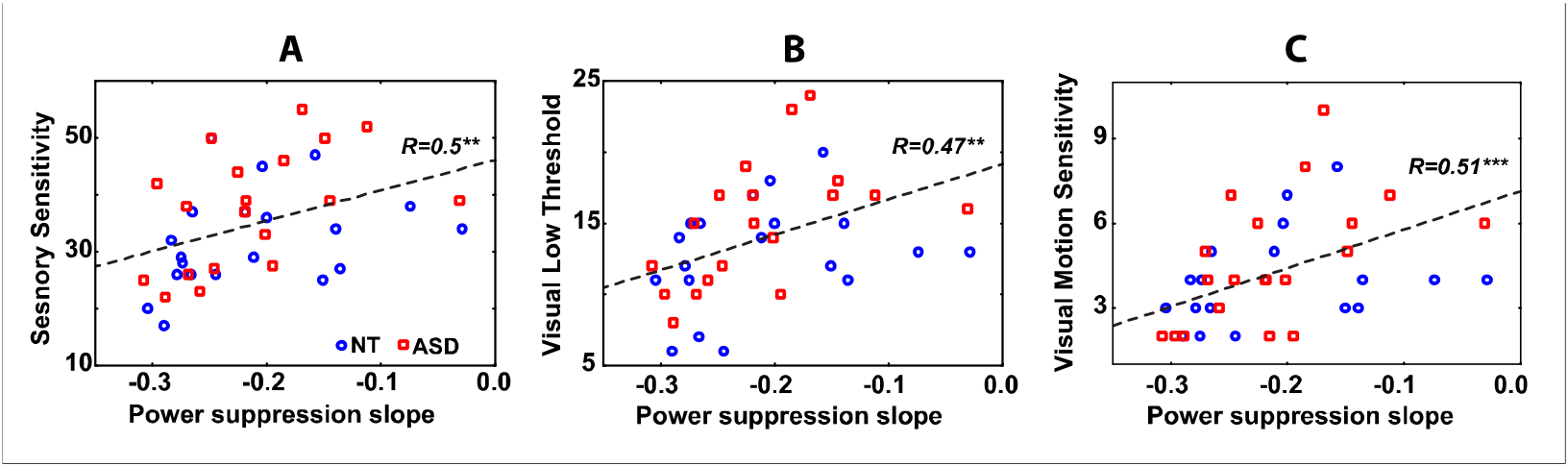
The relationship between gamma suppression slope and Sensory Sensitivity (A), Visual Low Threshold (B), and Visual Motion Sensitivity (C) in the combined sample of ASD (read squares) and NT (blue circles) individuals.

The GSS is a relational measure and its correlations with the sensory sensitivity could be predominantly driven by gamma responses at particular velocities of the visual motion. In accord with the ‘gain control’ hypothesis the variability in gamma response strength to the most intensive stimulation (‘fast’ visual motion) should make major contribution to the individual variation in sensory sensitivity. Indeed, as predicted, the generally elevated visual sensitivity/avoidance and, in particular, sensitivity to visual motion were associated with higher gamma responses to the fast motion (Tab. 3).

**Table 3.**
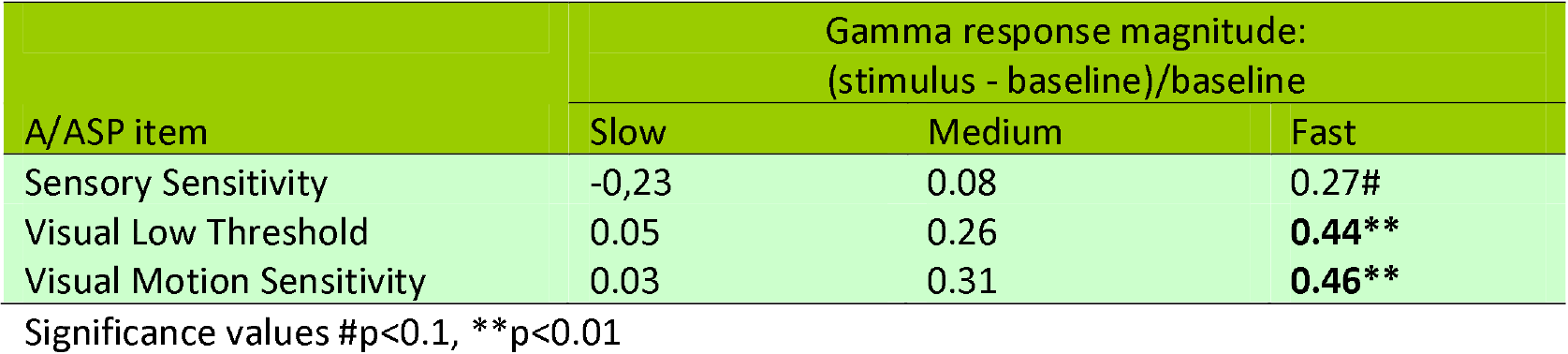
Spearman correlations between gamma response strength in the three velocity conditions and the A/ASP measures in the combined sample (NT+ASD).

Unlike the gamma suppression slope, neither the peak frequencies of gamma oscillations nor their velocity-related changes were related to the A/ASP measures of sensitivity (all p’s>0.05).

## Discussion

Our study shows that inter-individual variations in sensory sensitivity are strongly related to the capacity to modulate MEG gamma oscillations according to intensity of the visual input. We found that subjects who reported heightened sensory sensitivity were characterized by weakened suppression of the gamma response with increasing velocity of visual motion. This result indicates that the neural mechanisms underlying gamma suppression also modulate subjective reactivity to sensory events in the everyday life. Moreover, the similar pattern of findings in the NT and ASD individuals suggests that sensory hypersensitivity shares a common neural ground in people with autism and in general population.

In both NT and ASD subjects, the increase in velocity of the visual motion from 1.2 to 6°/s elicited a strong and reliable suppression of the visual gamma response accompanied by a substantial increase in gamma frequency for almost 15 Hz (Fig 3). These findings extend our previous results on the velocity-related changes of visual gamma response in NT subjects (Orekhova et al., 2015; Orekhova et al., 2018) and children with ASD (Stroganova et al., 2015) by replicating these findings in a group of adult ASD individuals.

Increasing motion velocity of full-contrast visual gratings up to 6.0°/s likely promotes excitation of interconnected excitatory and inhibitory neurons in the visual cortical areas (see (Orekhova et al., 2018) for discussion). According to the computational modeling results of Borgers and colleagues, increasing excitation of the I-neurons above some critical threshold leads to neuronal de-synchronization and thus suppression of gamma oscillations (Borgers and Kopell, 2005; Borgers and Walker, 2013; Cannon et al., 2014). This inhibitory-based physiological mechanism offers a reasonable explanation for the relative suppression of the induced gamma response at high stimulation intensities (visual motion at 3.6 and 6°/s, Fig. 3) in our study. Since gamma synchrony increases the impact of synaptic input from a neuronal group onto its postsynaptic targets (Fries, 2005, 2009, 2015; Ni et al., 2016; Vinck et al., 2013), the reduction of gamma power at high stimulation intensities may limit signal transmission between activated neural assemblies, thus protecting them from sensory-driven hyper-excitation. Hence, gamma suppression may reflect a suppressive gain-control mechanism, which affects sensory perception by reducing the impact of high-intensity stimulation. In a similar vein, a weaker suppression of visual gamma oscillations at the high motion velocities may be associated with a heightened behavioral sensitivity to the high-intensity visual stimulation.

To pursue this hypothesis, we introduced a measure that quantified the suppression of the gamma response with increasing stimulation intensity -- the ‘gamma suppression slope’ (GSS). As expected, a weaker negative slope (i.e. lower gamma suppression and less efficient homeostatic regulation of the E/I balance) correlated with a higher incidence of sensory noticing/discomfort and avoidant behaviors in both the NT and the ASD individuals (Table 2 and Fig.3). Presence of similar correlation patterns in the NT and ASD subjects suggests that variations in response gain control contribute similarly to individual differences in sensory sensitivity in the ASD group and the general population. The higher correlations in the ASD individuals can be explained by somewhat greater variability of their sensory sensitivity scores comparing to those in the NT subjects. The fact that the highly sensitive individuals displayed elevated gamma responses to the high-intensity stimuli, rather than reduced responses to those at lower intensities (Table 3), gives additional support for the suggested link between sensory hypersensitivity and inefficient down-regulation of excessive activation of the visual cortex. In general, these results confirm our hypothesis that the mechanisms leading to gamma suppression in response to strong sensory input also serve to protect the brain from hyper-excitation.

Most probably, the suppression of the gamma response at the highest stimulus velocity/temporal frequency used in our study (6°/s or 10Hz) is associated mainly with decrease in the efficacy of transient interactions between neural populations rather than with reduction of neuronal firing. Indeed, a recent study in monkeys showed that the 50-80 Hz power in local field potential (LFP) recordings reaches its maximum at lower temporal frequencies of visual motion than the neuronal spiking does (Salelkar et al., 2018). In particular, when the temporal frequency of visual motion increased from 2 to 8 Hz the LFP gamma suppression in monkeys was paralleled by an increase or absence of change in neuronal firing. Similarly, in humans, BOLD activation in visual cortical areas elicited by drifting visual gratings drops only after increasing temporal frequency beyond 9 Hz (Singh et al., 2000), suggesting high level of cortical activation at this temporal frequency. According to the model of Borgers and Kopell (2005), the asynchronous activity of the over-excited I-neurons, corresponding to the ‘no gamma state’, can suppress activity of the E-neurons. It is therefore likely that motion velocities/temporal frequencies yet higher than those applied in our study would result in a complete blockage of gamma oscillations paralleled by a decrease in E-neurons firing in the visual cortex.

Given that the homeostatic control of neural excitability may differ between cortical areas, we expected that the suppression of visual gamma response would be most closely related to behavioral sensitivity in the visual domain. Indeed, in case of the ‘Low Threshold’ A/ASP measures, the correlation with the GSS was highest for the visual modality (*Supplementary Table 2*). Correlations with the GSS in non-visual sensory or behavioral domains can be explained by presence of common neural factors affecting response gain control across sensory modalities, i.e. global variations in neural excitability, functional or structural connectivity, etc.

It is noteworthy that the majority of the ASD and NT participants in our study differed neither in regard to visual sensitivity (Table 1), nor gamma parameters (Fig. 2). These results suggest that the capacity to down-regulate growing neural excitation with increasing intensity of a sensory input was relatively preserved in the visual cortex in our high-functioning adult participants with ASD. Although an altered E/I balance is thought to be an important mechanism of ASD (Levin and Nelson, 2015; Rubenstein and Merzenich, 2003), it is possible that it is less affected (or better compensated for (Nelson and Valakh, 2015)) in the visual cortex than in other cortical areas. For example, Gaetz and colleagues (Gaetz et al., 2014) reported that the concentration of the inhibitory transmitter gamma-aminobutyric acid (GABA) was normal in the visual cortex of adolescents with ASD, whilst being significantly reduced in their auditory and motor cortices. Yet another possibility is that our participants with ASD represented a subgroup characterized by a relatively low prevalence of atypical sensory sensitivity, including that in the visual domain, compared to a more general ASD population. For example, in the study of Crane et al, the Sensory Sensitivity scores in adults with ASD were higher than in our study (Crane et al: 45.0; this study: 37.7), while the corresponding scores for the NT subjects were more similar (Crane et al: 33.8; this study: 31.2).

Recent advances in genetics and neuroscience clearly demonstrate that behavioral symptoms of ASD and other neurodevelopmental disorders may stem from cardinally different genetic and molecular etiologies that cause either increases or decreases in the E/I ratio (Lee et al., 2017; Nelson and Valakh, 2015). Given the heterogeneous nature of ASD (Gillberg, 2010; Jeste and Geschwind, 2014; Tordjman et al., 2018), the abnormal capacity to regulate the E/I balance in the visual cortex could characterize only a proportion of ASD individuals, as well as, e.g., patients with fragile X syndrome who are often hypersensitive to visual stimuli and are suggested to have elevated neural excitability of the visual cortex (Rigoulot et al., 2017; Schneider et al., 2009; Sinclair et al., 2017; Van der Molen et al., 2012). Considering the reliable correlation between sensory sensitivity and GSS (Fig. 5), one may predict that these individuals would demonstrate strongly reduced gamma suppression. In this respect, the GSS measure may provide a biomarker that can be used to select ASD subgroups according to a distinct neural phenotype – E/I imbalance in the visual cortex. Considering that many promising pharmacological agents tested or being testing in the ASD in clinical trials target the E/I balance, stratification of this clinical population according to the relevant neural deficits is important for selection of an individually appropriate treatment and tracking the treatment outcome.

Our study has several limitations. *First*, our participants were nearly exclusively males. An additional study is needed to generalize the results to females. *Second*, each of the experimental groups had a relatively small sample size, which stress the need for an independent replication study. Besides, as a group, our participants with ASD did not differ in their sensory responsiveness from the NT individuals, and it would be important in the future to investigate gamma suppression parameters in individuals characterized by excessive visual hypersensitivity. *Fourth*, our experimental paradigm was specifically aimed at testing the gain control in the visual cortex, and it’s unclear whether a similar modulation of cortical gamma responses by input intensity is present in other sensory modalities, e.g. auditory or tactile.

In conclusion, the modulation of gamma response power by intensity of visual input may give important information about the neural mechanisms that mitigate rising excitation and maintain E/I balance in the visual networks. Given the need for sensitive and objective measures of region-specific cortical excitability in different patient populations, this input-output relationship in gamma response strength offers a promising translational tool for clinical research. We suggest that the slope of the stimulus-response function of visual gamma oscillations may provide a tractable and accessible measure of the capacity to regulate the E/I balance in visual circuitry according to intensity of the visual input. It could be an especially appropriate measure in some groups of patients characterized by an elevated cortical excitability and high sensitivity to visual stimuli, such as patients with photo-sensitive epilepsy, migraine, and some forms of ASD. This non-invasive biomarker for unbalanced cortical excitability could also be used to select distinct sub-groups of patients within heterogeneous clinical populations (e.g., within ASD) and to track the impact of tailored pharmacological interventions in clinical trials.

## Materials and methods

### Participants

Twenty individuals (1 female) with ASD were included in the study, these were drawn from two study groups that have been described elsewhere (Davidsson et al., 2017; Helles et al., 2015). Briefly, 14 individuals had been assigned an ASD-diagnosis at three different occasions by structured clinical interviews. The remaining 5 individuals had been assigned an ASD-diagnosis at the Clinical Neuropsychiatry Centre in Gothenburg and then via a parental interview. One individual with ASD was recruited via advertisement, whose health journals were scrutinized and reviewed by a senior child and adolescent psychiatrist in order to verify the diagnosis. Nineteen ‘neuro-typical’ (NT) participants were recruited via advertisement. The NT participants underwent a brief screening focusing on neurological and psychiatric disorders in order to rule out psychopathology. To assess cognitive ability the Wechsler Adult Intelligence Scale, fourth edition (WAIS-IV) was used (for a few subjects in the ASD-group WAIS-III data was used). Individuals with an IQ below 80 were excluded. The NT and ASD group did not differ significantly in either age (ASD: 18.8-50.0 years, mean=31.1, SD=7.9; NT: 19.2-40.1 years, mean=27.3, SD=6.4; p>0.1) or general IQ (ASD: 78-140, mean=108.8, SD=15.7; NT: 96-135, mean=114.0, SD=11.5; p>0.2). The study has ethical approval from the regional ethical review board in Gothenburg (DNR: 552-14). Participants followed the informed consent procedure and were repeatedly given the option to discontinue their participation in the study.

### Assessment of sensory function

All the subjects filled in the Adolescent/Adult Sensory Profile questionnaire (A/ASP) (Brown and Dunn, 2002). This instrument combines information about sensory processing into four categories: ‘Low Registration’, ‘Sensation Seeking’, ‘Sensory Sensitivity’, and ‘Sensation Avoiding’. Here, we were interested in the Sensory Sensitivity scale that measures passive behavioral responses that characterize an individual’s sensitivity to environmental events, such as noticing behaviors, distractibility, and discomfort with sensory stimuli.

The A/ASP also allows assessment of the ‘Neurological Thresholds’ by combining the items across categories. Combined scores on the ‘Sensory Sensitivity’ and ‘Sensation Avoiding’ categories constitute the ‘Low Neurological Threshold’ (called below ‘Low Threshold’) that measures a person’s notice of, or annoyance with, sensory stimuli. This ‘Low Threshold’ category can be pooled for sensory modalities, as well as calculated separately for sensory/behavioral domains. Here we were mainly interested in the ‘Low Threshold’ for the visual modality.

Since we used moving visual stimuli in the present study, the velocity-related suppression of gamma response might most closely reflect subject’s sensitivity to the *moving* visual stimuli. Therefore, we also introduced the ‘Visual Motion Sensitivity’ scale by combined two A/ASP items that measured subject’s discomfort associated with intensive visual motion (i.e. Item 22: ‘ *I am bothered by unsteady or fast moving visual images in movies or TV’*; Item 25: ‘*I become bothered when I see lots of movement around me (for example, at a busy mall, parade, carnival)*’).

### Experimental task

To measure gamma, we applied an experimental paradigm that has been shown to induce reliable MEG gamma responses in the visual cortex in our previous studies (Orekhova et al., 2015; Orekhova et al., 2018; Stroganova et al., 2015). The stimuli were generated using Presentation software (Neurobehavioral Systems Inc., USA) and presented using a FL35 LED DPL gamma-corrected projector with 1920×1080 screen resolution and 120 Hz refresh rate. They consisted of black and white sinusoidally modulated annular gratings with a spatial frequency of 1.66 cycles per degree of visual angle and an outer diameter of 18 degrees of visual angle. The gratings appeared in the center of a screen over a black background and drifted to the central point at velocities of 1.2, 3.6, or 6.0°/s, (which approximately corresponds to temporal frequencies of 2, 6, and 10 Hz, respectively); hereafter, we respectively refer to these three velocities as ‘slow’, ‘medium’, and ‘fast’. Each trial began with the presentation of a white fixation cross in the center of the display over a black background for 1200 ms that was followed by the moving grating that persisted for 1200-3000 ms and then stopped. The participants were instructed to respond to the termination of motion with a button press. If no response occurred within 1 second, the grating was substituted by a discouraging message “too late!” that remained on the screen for 2000 ms, after which a new trial began. Error trials (misses or responses that occurred <150 ms after the stop) were excluded from the MEG analysis. Stimuli were presented in three experimental blocks in a random order resulting in 90 repetitions of each stimulus type. The luminance of the screen measured at the position of the observer’s eyes was 53 Lux during the stimulation and 2.5 Lux during the inter-stimulus interval. Short (3-6 s) animated cartoon characters were presented randomly between every 2-5 stimuli to increase vigilance and minimize fatigue.

### MEG recording

MEG was recorded at the NatMEG Centre (The Swedish National Facility for Magnetoencephalography, KarolinskaInstituted Stockholm) using 306-channel system (ElektaNeuromag TRIUX). The data was recorded with a band-pass filter of 0.1-330 Hz, digitized at 1000 Hz, and stored for off-line analysis. The subjects’ head position during MEG recordings was continuously monitored.

### MRI recording

Structural brain MRIs (1 mm^3^ T1-weighted) were obtained for all participants and used for source reconstruction.

### MEG data preprocessing

The data was first de-noised using the Temporal Signal-Space Separation (tSSS) method (Taulu and Hari, 2009) and adjusted to a common head position. The de-noised data was filtered between 1 and 145 Hz and resampled at 500 Hz. Independent component analysis (ICA) was used for correction of biological artifacts. The data was then epoched (−1 to 1.2 s relative to the stimulus onset) and checked for the presence of residual artifacts. After rejection of the artifact-contaminated epochs and error trials, the average number of the ‘good’ epochs for the ‘slow’, ‘medium’ and ‘fast’ conditions was 79.5, 79.1, 77.1 in the NT, and 75.4, 75.9, 76.9 in the ASD, groups. No group differences in the numbers of valid trials were found (all p’s>0.3). Details for the preprocessing are given in the *Supplementary Methods*.

### MEG source analysis

Source localization was performed using linearly constructed minimum variance (LCMV) beamformer implemented in the FieldTrip M/EEG toolbox (Oostenveld et al., 2011) (see *Supplementary Methods* for details). For each subject and experimental condition, the weighted peak gamma power and frequency were then calculated for the average spectrum of the virtual sensors in 26 voxels closest to and including the ‘maximally modulated voxel’ (the voxel in the visual cortex with highest increase of 45-90 Hz power during stimulation). A frequency range of interest was defined as those frequencies where the (stimulus-baseline)/baseline power ratio exceeded 2/3 of the maximum for the particular subject and condition. The gamma peak frequency was calculated as the center of gravity, whereas the peak power was the average power, over that range. For each subject/condition, we also calculated probabilities of the post-stimulus increase in the 45-90 Hz gamma power in the selection of the 26 voxels. The individual peak gamma frequencies were analyzed only if the probability of the gamma power increase during stimulation relative to pre-stimulus period was significant at p<0.0001.

In order to quantify the suppression of the gamma response power with increasing visual motion velocity, we introduced the ‘gamma suppression slope’ index. We calculated the coefficient of regression of the weighted gamma power to velocity using the ‘fitlm’ Matlab function: fitlm (x,y,‘y^~^x1−1’); where x=[1.2, 3.6, 6.0], y=[0, POW_medium_/POW_slow_−l, POW_fast_/POW_slow_−l] and ‘y^~^x1−1’ sets the intercept of the regression line to zero. The resulting regression coefficient b is equal to zero in the case of a constant response power in the three experimental conditions (i.e. ‘no suppression’) and is proportionally more negative in case of stronger suppression of the gamma response with increasing motion velocity (Fig. 4).

The suppression of gamma response with increasing velocity can be reliably estimated only if a reliable response is observed in the slow velocity condition (i.e. the condition that induces a greater gamma response than the ‘medium’ and ‘fast’ conditions (Orekhova et al., 2015)). Therefore, we estimated the power suppression slope only if the probability of the post-stimulus gamma increase in the ‘slow’ condition relative to prestimulus baseline was high (p<0.0001); this lead to exclusion of one ASD participant from the correlation analysis.

### Statistical analysis

T-test was applied to analyze group differences in the A/ASP measures. Since the distribution of some MEG parameters failed a normality test, we used Spearman coefficients for correlation analyses. The repeated measures ANOVA was initially used to test for the effects of Condition, Group, and Group × Condition interaction on the gamma parameters, and the post-hoc comparisons were performed using the nonparametric Mann-Whitney U-test.

## Acknowledgements

This work was financed by the Torsten Soderberg Foundation (M240/13 to CG) and the Russian Scientific Foundation (14-35-00060 to TS). Authors JFS and BR are supported by the Knut and Alice Wallenberg Foundation (grant 2014.0102), the Swedish Research Council (grant 621-2012-3673), and the Swedish Childhood Cancer Foundation (grant MT2014-0007). The author NH was supported by the LifeWatch Foundation. The author TS was supported by the Charity Foundation “Way Out”. We heartily thank all participants for their participation in this study.

## Financial Disclosures

The authors report no financial interests or potential conflicts of interest.

